# Oscillating bacterial expression states generate herd immunity to viral infection

**DOI:** 10.1101/244814

**Authors:** Christopher J. R. Turkington, Andrey Morozov, Martha R. J. Clokie, Christopher D. Bayliss

## Abstract

Hypermutable loci are widespread in bacteria as mechanisms for rapid generation of phenotypic diversity, enabling individual populations to survive fluctuating, often antagonistic, selection pressures. As observed for adaptive immunity, hypermutation may facilitate survival of multiple, spatially-separated bacterial populations. We developed an ‘oscillating prey assay’ to examine bacteriophage (phage) spread through populations of *Haemophilus influenzae* whose phage receptor gene, *lic2A*, is switched ‘ON’ and ‘OFF’ by mutations in a hypermutable tetranucleotide repeat tract. Phage extinction was frequently observed when the proportion of phage-resistant sub-populations exceeded 34%. *In silico* modelling indicated that phage extinction was interdependent on phage loss during transfer between populations and the frequency of resistant populations. In a fixed-area oscillating prey assay, heterogeneity in phage resistance was observed to generate vast differences in phage densities across multiple bacterial populations resulting in protective quarantining of some populations from phage attack. We conclude that phase-variable hypermutable loci produce bacterial ‘herd immunity’ with resistant intermediary-populations acting as a barricade to reduce the viral load faced by phage-sensitive sub-populations. This paradigm of meta-population protection is applicable to evolution of hypermutable loci in multiple bacteria-phage and host-pathogen interactions.

**Importance:** Herd immunity is a survival strategy wherein populations are protected against invading pathogens by resistant individuals within the population acting as a barrier to spread of the infectious agent. Although, this concept is normally only applied to higher eukaryotes, prokaryotic organisms also face invasion by infectious agents, such as bacterial viruses, bacteriophage (phage). Here we use novel experimental approaches and mathematical modelling, to show that bacteria exhibit a form of herd immunity through stochastically generated resistant variants acting as barricades to phage predation of sensitive cells. With hypermutable loci found in many prokaryotic systems, this phenomenon may be widely applicable to phage-bacteria interactions and could even impact phage-driven evolution in bacteria.

## Introduction

Hypermutable loci as mediators of survival against constantly fluctuating selection pressures are a predictable outcome of the evolution of evolvability as stated in the Red Queen hypothesis (1, 2). Fluctuating selection pressures are regularly faced by bacteria during persistence in human hosts, where bacteria adhere to host surfaces while contending with varying nutrient concentrations, frequent exposure to immune effectors and predation by bacteriophages. These fluctuations often select for and against opposing gene expression states of single loci leading to evolution of localised hypermutable mechanisms that produce frequent switches in single-gene expression states.

One class of hypermutable loci facilitate survival of conflicting selective pressures by pre-emptive, frequent, and reversible ‘ON/OFF’ generation of adaptive variants, in a process known as ‘phase variation’ (PV; 2–5). A major mechanism of PV involves increases and decreases in identical, tandemly-arranged DNA repeats (microsatellites) by slipped strand mispairing during DNA replication (3, 5, 6). The obligate human respiratory commensal and pathogen *H. influenzae* contains an expansive array of repeat driven phase variable loci (7–13). Several of these loci are required for addition of sugar molecules onto the surface-exposed outer-core of the lipooligosaccharide (LOS) (5, 7). PV of the UDP-galactose-LOS-galactosyltransferase encoding gene, *lic2A*, is mediated by a 5’CAAT repetitive tract present in the open-reading frame (14). Phage HP1c1 attaches to an LOS epitope of *H. influenzae* strain Rd that contains the galactose sugar added by *lic2A* (15). PV of *lic2A* causes switching between phage sensitive (*lic2A* ON) and phage resistant states (*lic2A* OFF) (15, 16). Partial resistance to phage HP1c1 in strain Rd is also mediated by PV of a Type I restriction-modification (RM) system. PV of surface receptors and RM systems generates resistance to phage infection in several bacterial species (17–22).

While phage-receptor PV prevents viral propagation in individual populations, the frequency and distribution of resistant variants within larger meta-populations may also impose inhibitory effects on phage spread through multiple spatially-linked sub-populations. In order to explore this potential benefit, we examined how diversity in phage resistance/sensitivity phenotypes generated by one hypermutable locus, *lic2A*, alters spread of phage HP1c1.

## RESULTS

### Low numbers of resistant bacterial populations significantly restrict phage spread and densities

PV can generate high levels of ON and OFF variants of single genes within individual populations but also has the potential to generate population-to-population variation within a meta-population. For example, *lic2A*-positive *H. influenzae* colonies on agar plates will have most cells in a *lic2A* ON expression state but will also contain small, but significant, numbers of *lic2A* OFF variants. Similarly, an *H. influenzae* meta-population may consist of multiple sub-populations with one fraction being *lic2A*-positive and another fraction being *lic2A*-negative. Such population-to-population variation is observed for *H. influenzae* colonies on agar plates and for *H. influenzae* populations isolated from artificially-inoculated animals and asymptomatic human carriers (23–27). Proportions of ON and OFF sub-populations in a meta-population will depend on levels of selection for each expression state and on ‘founder’ effects that influence the starting state of each sub-population. Thus, any phage invading a bacterial meta-population where there is PV of the receptor must contend with highly variable distributions of resistance and sensitive sub-populations.

To simulate the effect of phage-receptor PV on spread of phage through bacterial meta-populations we developed the ‘oscillating prey assay’ (Fig. 1). This assay involves continual cycling of phage through *H. influenzae* strain Rd cultures with either a majority *lic2A* ON or OFF phenotype. Each cycle allows for one round of phage replication after which the phage-containing supernatant is recovered, filtered, and transferred to a new culture of either *lic2A* ON or OFF *H. influenzae* cells. During transfer, the supernatant is subject to a 10-fold dilution. This arbitrary dilution factor simulates loss of phage during population-to-population transmission.

**Fig. 1.**
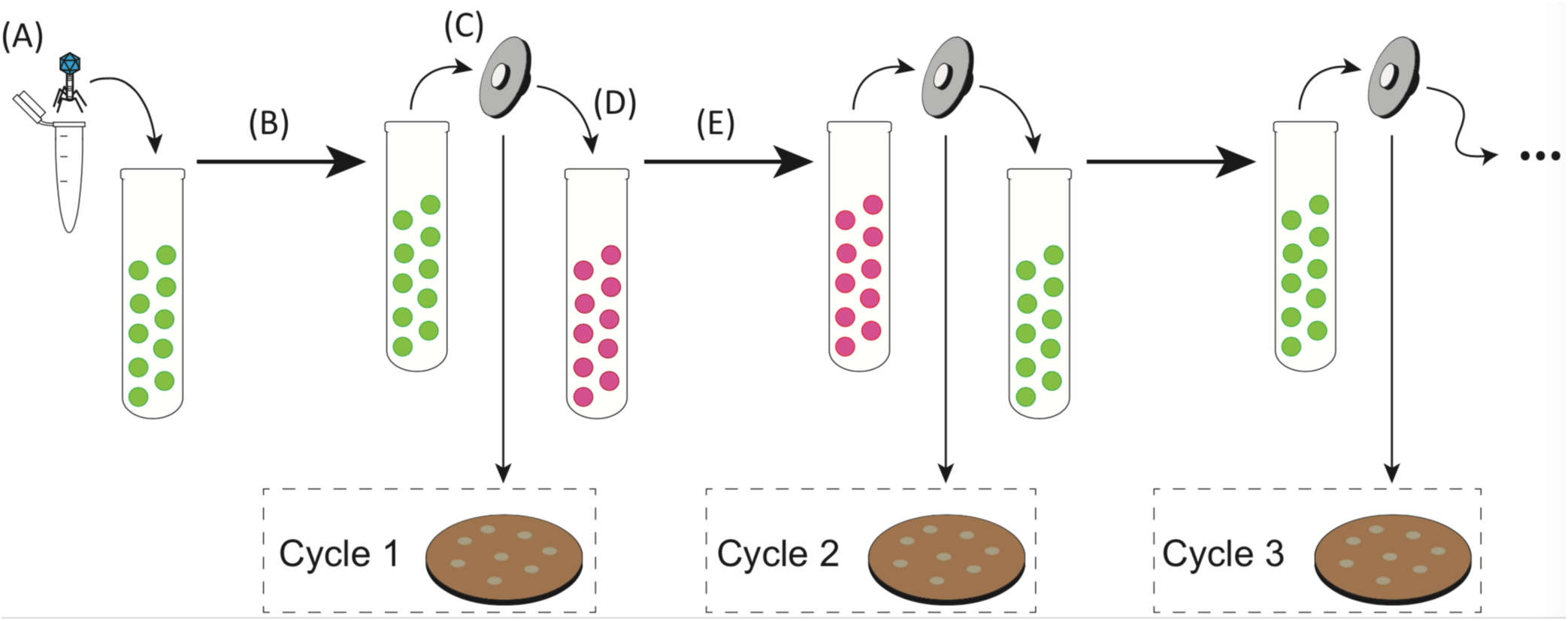
Graphic representation of the oscillating prey assay. This diagram illustrates the methodology for experimental simulation of phage HP1c1 transmission through a 50 % ON meta-population structure. Green circles represent a phage-sensitive Rd 30S culture (*lic2A* ON) while pink circles represent a phage-resistant Rd 30R culture (*lic2A* OFF). (A) Phage are added to an exponential phase culture of the bacterial phase variant with an OD_600_ of 0.01 to a final MOI of 0.01 (i.e. ∼1 × 10^6^ PFU/mL). (B) Incubation of phage with host at 37°C for 50 minutes to allow for one viral replicative cycle. (C) Filtration of the phage-bacteria mix through a 0.22 µm filter and titration to determine the phage density at the end of the cycle. (D) Transfer of 600 µL of filtered culture-suspension to 5.4 mL of fresh culture of the appropriate phase variant (in this example Rd 30R). (E) Incubation of this new culture at 37°C for 50 minutes beyond which the process is repeated until a total of 20 cycles have been completed.

Six population structures were examined in the oscillating prey assay: [1] 100 % ON (HP1c1 cycled only through *lic2A* ON populations; S100), [2] 66 % ON (2:1 *lic2A* ON:OFF; S66), [3] 50 % ON (1:1 *lic2A* ON:OFF, starting with an ON culture; S50), [4] 50 % OFF (1:1 *lic2A* ON:OFF, starting with an OFF culture; R50), [5] 66 % OFF (1:2 *lic2A* ON:OFF; R66), and [6] 100 % OFF (HP1c1 cycled only through *lic2A* OFF populations; R100). Survival and propagation of phage was dependent on the proportion of phage-resistant sub-populations in each series of 20 cycles (Fig. 2). Survival of phage to the final cycle was only observed when the proportion of phage-resistance populations was ≤ 34% (i.e. the S66 and S100 populations; Fig. 2). Extinction events occurred within 5 to 16 cycles in all other heterogeneous and homogeneous populations at a rate that was dependent on the proportion of resistant sub-populations. Despite survival of phage through to cycle 20 in the S66 population series, phage densities were significantly decreased by this cycling regime (Fig. 2; paired *t*-test: *t* = 4.97, *P <* 0.05; mean ± SEM PFU/mL values for phage densities at cycle 0 = 7.16 ± 0.26 × 10^5^ and cycle 20 = 2.82 ± 0.9 × 10^4^), indicating that further passages with a similar pattern would have resulted in phage extinction. Contrastingly, phage density increased when all sub-populations were phage sensitive (S100), with phage densities plateauing after ∼11 cycles. Thus, both phage survival and density were limited by the frequency of encounters with phage-resistant *lic2A* OFF phase variants during passage through a linear series of sub-populations.

**Fig. 2.**
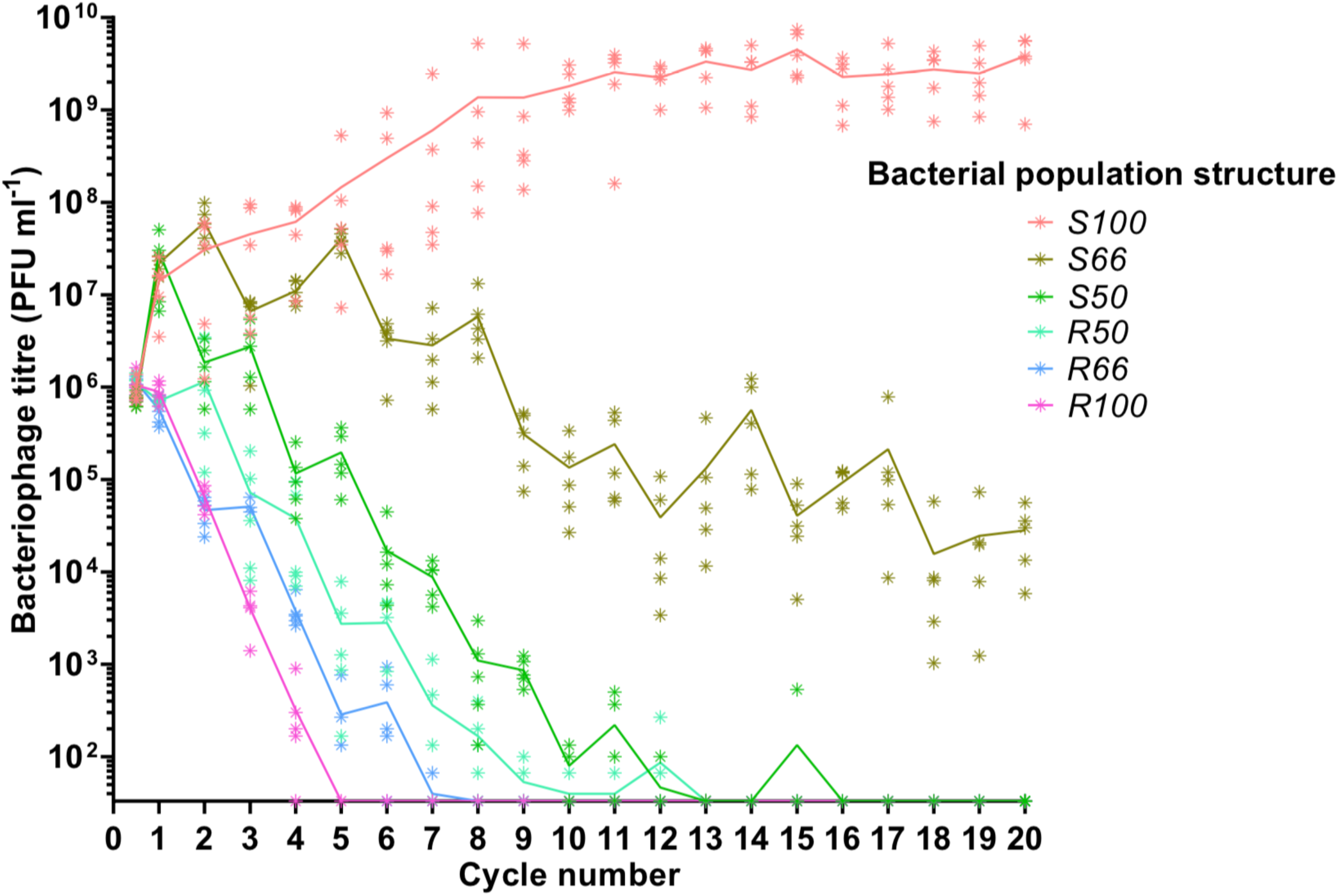
Oscillating prey assay for phage HP1c1 infections of *H. influenzae* strain Rd populations with varying population structures for *lic2A* expression. Each line represents cycling of the phage through a defined series of cultures of two *H. influenzae* strain Rd phase variants, namely Rd30S (S; *lic2A* ON; phage sensitive) and Rd 30R (R; *lic2* OFF; phage resistant). The population structures are indicated in the legend (e.g. S100, all S; S66, S-S-R-S-S-R; S50, S-R-S-R; etc.). The phage HP1c1 concentration was determined at the end of each cycle. Circles represent the phage titre observed from each of five biological replicates, with the line showing the mean of these replicates.

### Detection of combinatorial effects of population structure and dilution rate on phage extinction using an *in-silico* model of phage spread

Our experimental data indicated that meta-population structure had a major impact on phage survival and spread. In order to explore a wider range of linear and non-linear cycling patterns and the effects of dilution rate, we developed a mathematical model of the oscillating prey assay. This model utilized key phage parameters for adsorption rate, replication time, burst size and stability of phage HP1c1 (see Fig. S1). The number of oscillations was extended to 105 phage replicative cycles, while cycling patterns were randomised for each specific overall proportion of *lic2A* ON/OFF sub-populations. The mathematical model produced comparable findings to the experimental setting (Fig. S2). Multiple runs of this model exhibited stochastic variation in phage densities and extinction events as observed in the experimental model but with extinction events occurring over a wider band of replication cycles (Fig. 3A-B). This model demonstrates that random patterns of ON/OFF expression states for a phage receptor can limit phage spread.

**Fig. 3.**
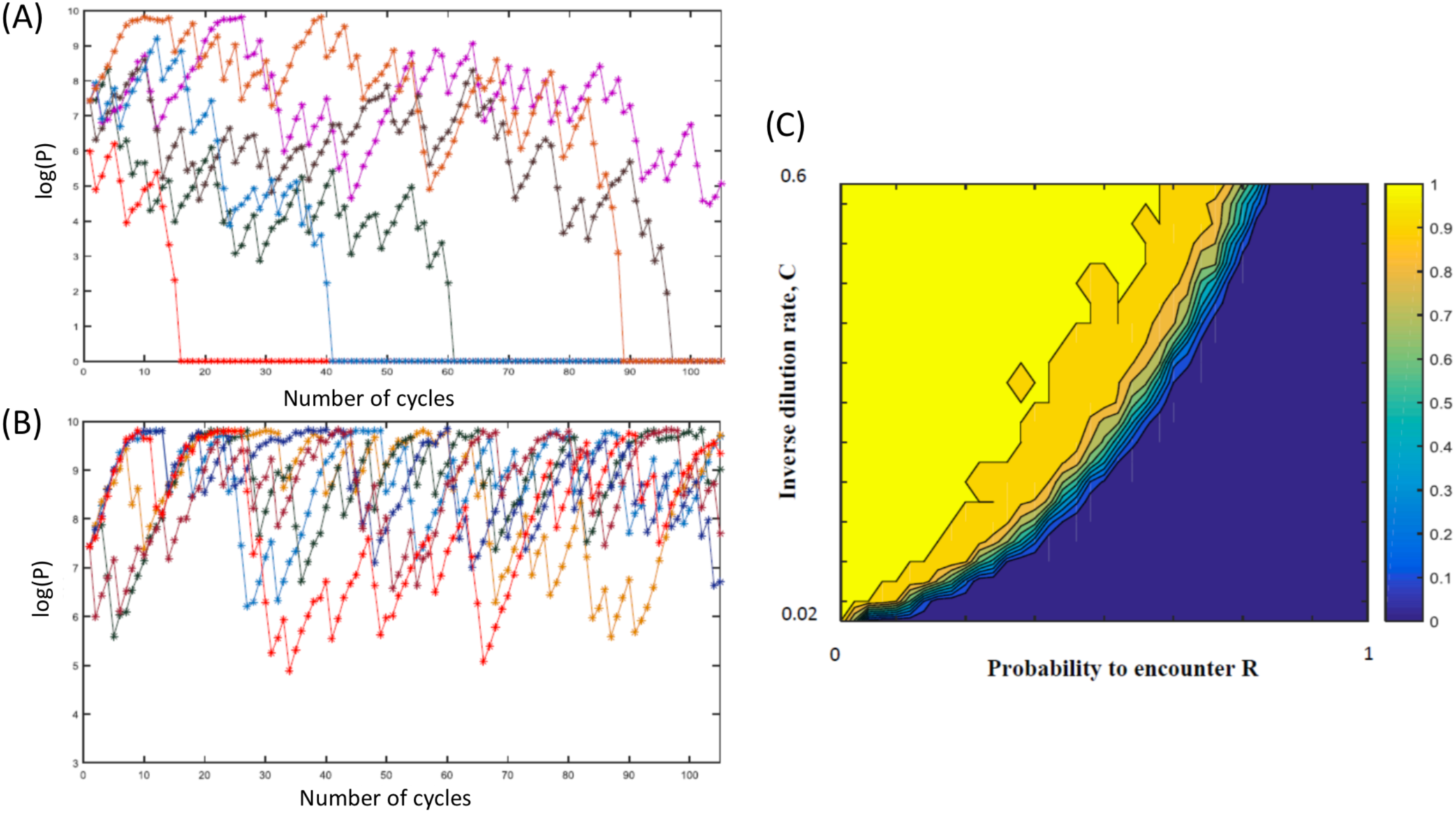
Mathematical model of the impact of sub-population phage resistance/sensitivity composition and dilution rate on phage extinction events. This model simulates transmission of phage HP1c1 through meta-populations of *H. influenzae* strain Rd comprising either phage-sensitive (S, *lic2A* ON) or phage-resistant (R, *lic2A* ON) populations for 105 cycles. In these simulations, the order with which phage met either S or R populations was random but dependent on the probability of encountering a resistant population (*R*). Note that the phage population is classified as extinct if the phage titre falls below the extinction threshold (*P*_*0*_ = 100) with the density being set to 0 for all remaining subsequent cycles. Panels A and B show examples of model outputs, which are quantified as phage densities log(P). These graphs show six iterations of meta-populations comprising either 75% (i.e. *R* = 0.25; panel A) or 85 % (i.e. *R* = 0.15; panel B) phage-sensitive populations. Panel (C) shows the mean proportion of phage extinction events occurring in 200 lineages for 200 combinations of probabilities to encounter R (i.e. 1 = 100 % resistant, 0 = 0 % resistant) and rates of phage loss during transmission after each cycle (inverse dilution rate, C; 0.02 = 2 % of phage carried through in each transfer). Colour correlations are shown on the bar to the right of the graph with 1 representing no extinction events while 0 is extinction in all lineages.

Multiple simulations (n = 200) were performed for each combination of R (the percentage of phage-resistant *lic2A* OFF sub-populations) and C (the inverse of the dilution coefficient). Average phage densities were measured for all cycles and runs of each combination of R and C, and phage extinction was defined to occur whenever the density fell below 100 PFU/mL. Phage extinction was always observed when the number of resistant states exceeded 70% of sub-populations, even at a low dilution rate of 1 in 2 (C=0.5; Fig. 3C). Similarly, when phage loss was ≥ 98 % at each transfer (i.e. C=0.02), phage extinction was observed for all meta-population structures except those consisting of >95% phage-sensitive sub-populations (Fig. 3C). Between these extremes, there was an accelerating trade-off between R and C with respect to phage extinction and average phage density; decreases in dilution rate were countered by a high prevalence of resistant sub-populations enabling bacterial populations to survive even when phage dispersal was low (Fig. 3C). This observation suggests that on-going evolution of localised hypermutability for a surface-exposed bacterial epitope could be tuned to the prevalence and density of phages capable of using the phase-variable epitope as a receptor for surface attachment.

### Localised hypermutation-driven herd-immunity produces regional variations in phage densities within bacterial meta-populations

Both our experimental and *in silico* oscillating prey assays demonstrated that phage spread was influenced by the linear pattern of phase-variant sub-populations. However, the distribution of phase variants across a surface (e.g. microcolonies on upper respiratory tract surfaces for *H. influenzae*) is anticipated to be random and hence to result in spatial effects on phage spread. Indeed, spatially structured environments are known to restrict phage spread (28). Spatial structuring was explored by examining phage transmission across meta-populations of fixed dimensions but with varying proportions of each *lic2A* expression state (Fig. S3). This fixed-area oscillating prey assay was initiated by inoculating phage into one well of a 96-well plate and then expanding outward by seeding each subsequent replicative cycle into neighbouring wells (see Fig. S4).

Passage of HP1c1 through heterogeneous populations of the fixed-area oscillating prey assay resulted in uneven phage densities across the meta-populations consisting of 50-66% phage-sensitive populations (Fig. 4B-4D). Conversely, homogenous densities were observed for high (Fig. 4A) or low (Fig. 4E and 4F) phage-sensitive distributions. When 66% of populations were phage-sensitive (Fig. 4B), densities ranged from 10^3^ to 10^10^ PFU/mL, whereas densities always exceeded 10^8^ PFU/mL if all population were sensitive (Fig. 4A). Phage densities in specific regions of heterogeneous populations were dependent on the direction of propagation with passage through multiple resistant or sensitive sub-populations resulting in low or high phage densities, respectively. This model shows how spatial meta-population heterogeneity could prevent equal dissemination of phage through host populations with phase-variable phage-receptors and aid survival of phage-sensitive sub-populations whose phenotypes may be beneficial for bacterial survival against other selective pressures.

**Fig. 4.**
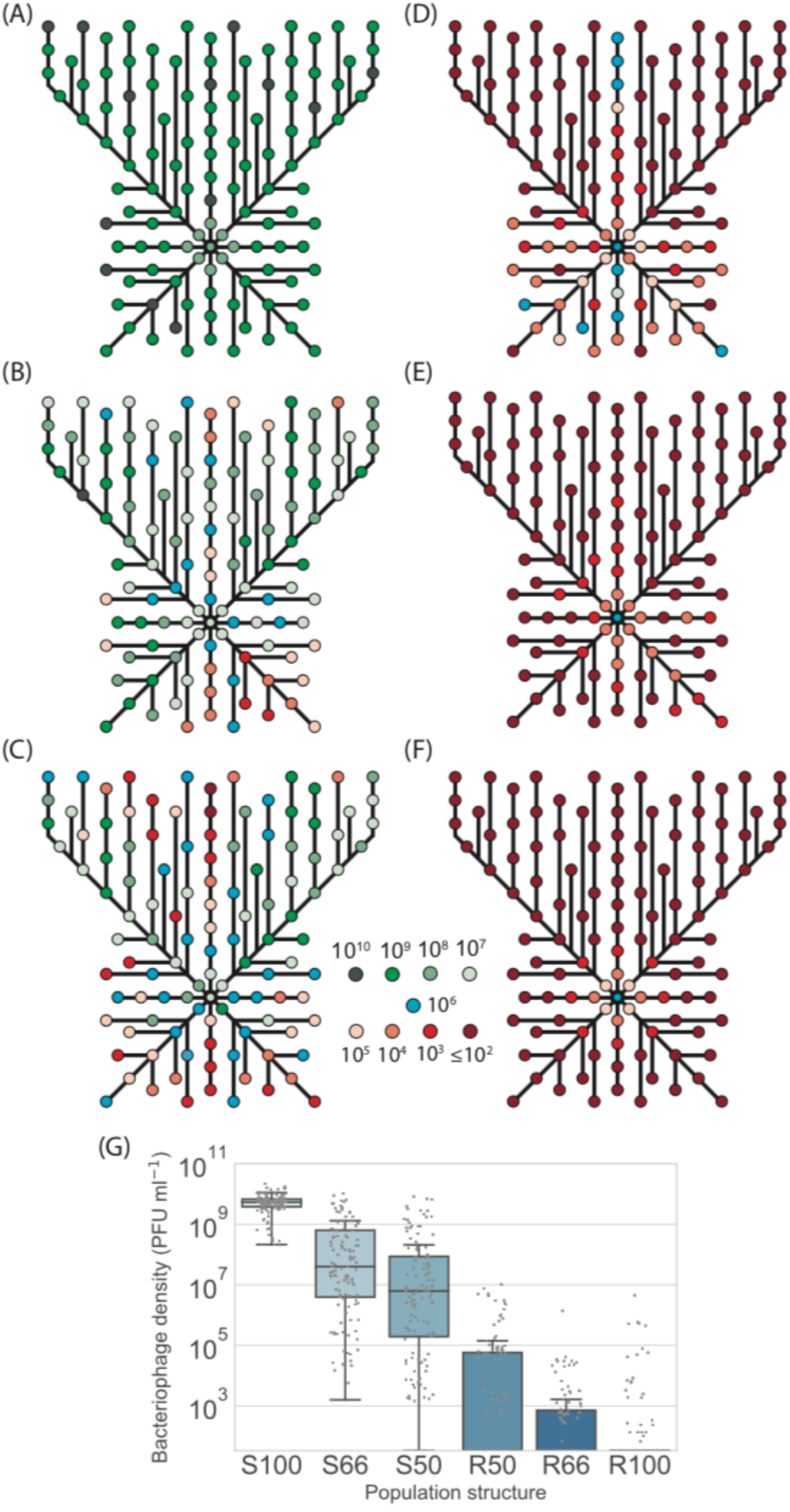
Illustration of phage survival in spatially-structured sub-populations of *lic2A* phase variants of *H. influenzae*. Phage HP1c1 was passaged through a two-dimensional array of phage-resistant (R) and phage-sensitive (S) populations. The proportion of phage sensitive sub-populations (*lic2A* ON) in each fixed area structure were as follows: (A) 100 % *lic2A* ON (S100); (B) 66 % *lic2A* ON (S66); (C) 50 % *lic2A* ON (S50); (D) 50 % *lic2A* ON (R50); (E) 33 % *lic2A* ON (R66); and (F) 0 % *lic2A* ON (R100). Each node represents a well in which phage density was measured. The colour of each node indicates the concentration of bacteriophage detected in a specific well. Lines indicate the route taken starting from a central initiator well with line length being proportional to the number of cycles between each node. All central initiator wells were inoculated with 10^6^ PFU/mL. (G) shows a box-plot of the distribution of phage densities for each tested well of the fixed-area oscillating prey assays. Densities of phage (PFU/mL) obtained from each test well are represented by a dot for the six distributions of *lic2A* phase variants (x-axis). The boxed area indicates the first to third quartile, the line is the median of all points, and whiskers represent 1.5x the interquartile range. Due to the nature of the small-drop plating methodology employed for phage enumeration the minimum detection threshold at any time point is 3.3×10^1^ PFU/mL.

### Observed PV rates generate populations with phase-variant proportions capable of herd-immunity

The fixed-area oscillating prey assay outputs showed how phage extinction was dependent on the proportion of phage-resistant sub-populations. *H. influenzae* normally resides in the upper respiratory tract where selection is likely to act on both PV states of a locus. Thus, the proportion of resistant populations depends on both selection strength for/against the phage-resistance phenotype, and the ON/OFF switching rate. Switching rates of *H. influenzae* phase-variable genes are malleable due to changes in repeat number and can evolve in response to alternating selection pressures (28, 29).

A mathematical model was developed to examine the impact of different switching rates and immune selection on the proportions of *lic2A* expression states (Fig. 5 and Supplementary Data S1). Dixon *et al.* (30) found that the *lic2A* ON-to-OFF (S-to-R; where R and S are the phage-resistant and phage-sensitive states, respectively) switching rate was 1.7-fold higher than the *lic2A* OFF-to-ON (R-to-S) switching rate. The mathematical model assumed that these proportions were maintained for three 10-fold differences in overall mutation rate representing low, intermediate and high repeats numbers (Fig. 5A-5C). In the absence of any selective difference between the S and R states (*m*=1), a high, steady-state proportion (>63%) of R, was observed for all mutation rates but with minor differences in the rate of approach to the steady state and absolute amounts of R. The *lic2A* OFF state (i.e. R) is known to be more immune sensitive than the ON state (i.e. S), we therefore imposed a selection against R. Even with strong selection (m=0.99), high levels of R were maintained by medium to high switching rates (Fig. 5A and 5C, respectively). In contrast, when switching rates were low (Fig. 5B), even weak selection (m=0.999) drives resistant variants to <1% (Fig. 5B). Our other *in silico* models indicated that phage spread was inhibited when the probability of encountering R variants was between 10-70% for dilution rates of 0.02 to 0.6. Thus, the immune model shows that observed repeat numbers and switching rates for the *lic2A* gene of *H. influenzae* strains can maintain sufficient phage-resistant variants for restricting phage spread even when there is immune selection against this state.

**Fig. 5.**
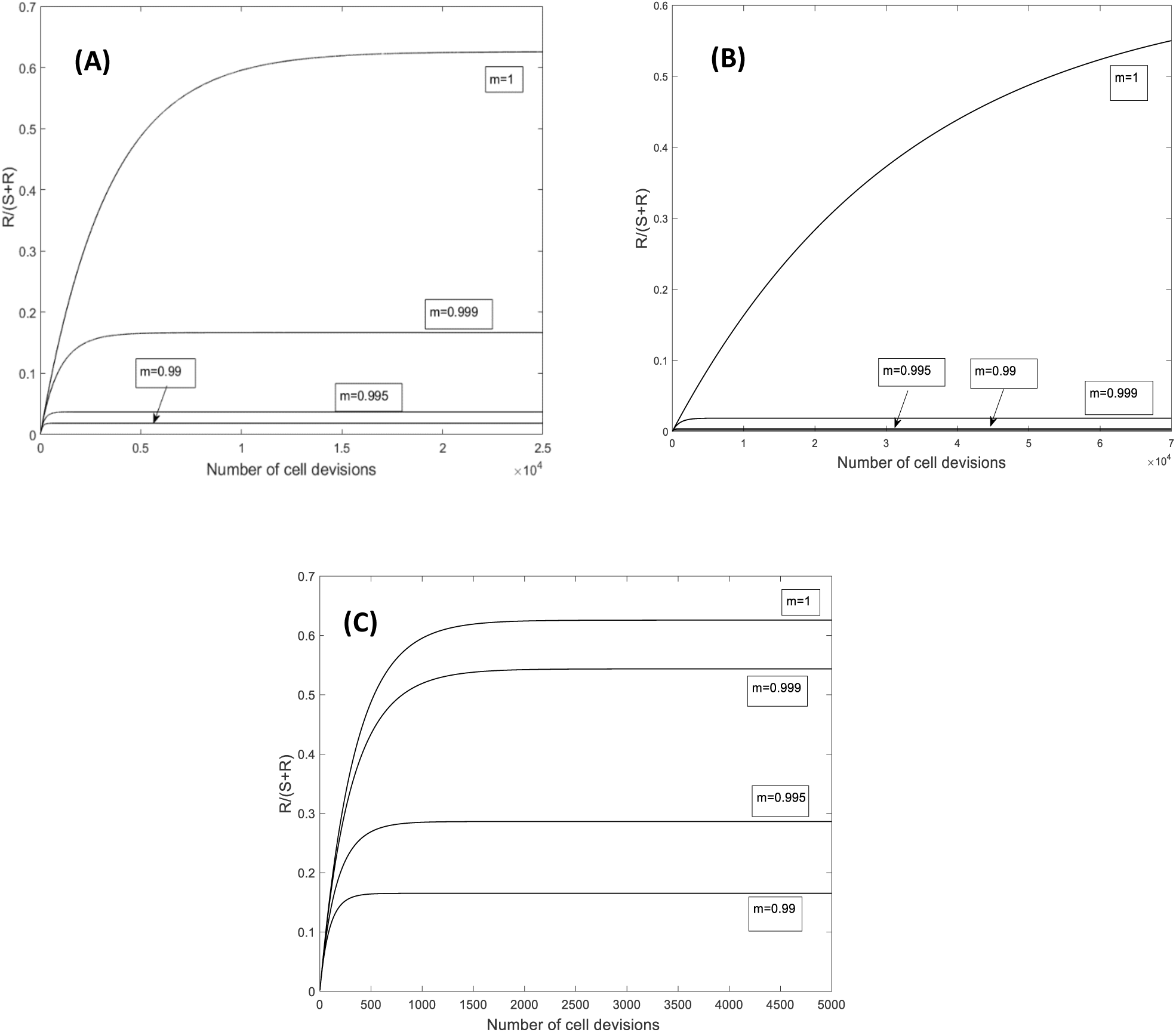
Model of the temporal fluctuations in the relative amounts of phage-sensitive and phage-resistant phase variants for a range of PV rates and selection pressures. PV of the *lic2A* gene results in switching between phage-sensitive (S, *lic2A* ON) and phage-resistant (R, *lic2A* OFF) phase variants. The *lic2A* ON state is however known to mediate serum resistance (see text). This model examines how the rate of *lic2A* PV (*β*, ON-to-OFF switching; *α*, OFF-to-ON switching; note that switching rates were obtained from Dixon *et al*. [30]) and the degree of selection (*m*) against the *lic2A* OFF (i.e. R, the phage-resistant state) expression state influences the relative amounts of the R and S states in a population. All panels show changes in the proportion of R. Three different switching rates were examined: *β* =1.89×10^−4^, *α* = 1.13×10^−4^ (A); *β* =1.89×10^−5^, *α* = 1.13×10^−5^ (B); *β* =1.89×10^−3^, *α* = 1.13×10^−3^ (C).

## DISCUSSION

Hypermutable loci have well-documented roles in facilitating survival of individual bacterial populations against phage predation through generation of phage-resistant cells. An unexplored concept is that hypermutation driven heterogeneity in phage resistance across the wider population also facilitates bacterial survival of phage predation.

### Phase-variable loci can generate herd immunity in bacterial meta-populations

The concept of herd immunity was derived to explain the protection of susceptible individuals in populations with high levels of immunity to an infectious agent as observed for measles in Baltimore (31). Ordinarily applied to naturally or vaccine-acquired immunity to infectious agents in human populations (32), we show, herein, that repeat-mediated PV of a phage-receptor provides a form of ‘bacterial herd-immunity’ at the population level (Fig. 6). Thus, phage spread is retarded by resistant sub-populations creating barriers between the phage and phage-sensitive bacterial sub-populations as shown in our one- and multidirectional experimental models (Fig. 2 and Fig. 5), and an *in silico* model with randomized patterns of phage sensitivity (Fig. 3). Key features of these models were that: the chance of phage survival was inversely proportional to the linear pattern of resistance faced by the phage population; random distributions of phage-resistant/sensitive populations result in large variations in viral numbers across a meta-population with some regions being completely free of phage; and dilution during transfer of phage between populations modulates the number of phage-resistant populations required to retard phage spread. Thus, localised hypermutation of a phage-receptor generates herd-immunity whereby phage-sensitive sub-populations can be maintained at high levels within bacterial meta-populations.

**Fig 6.**
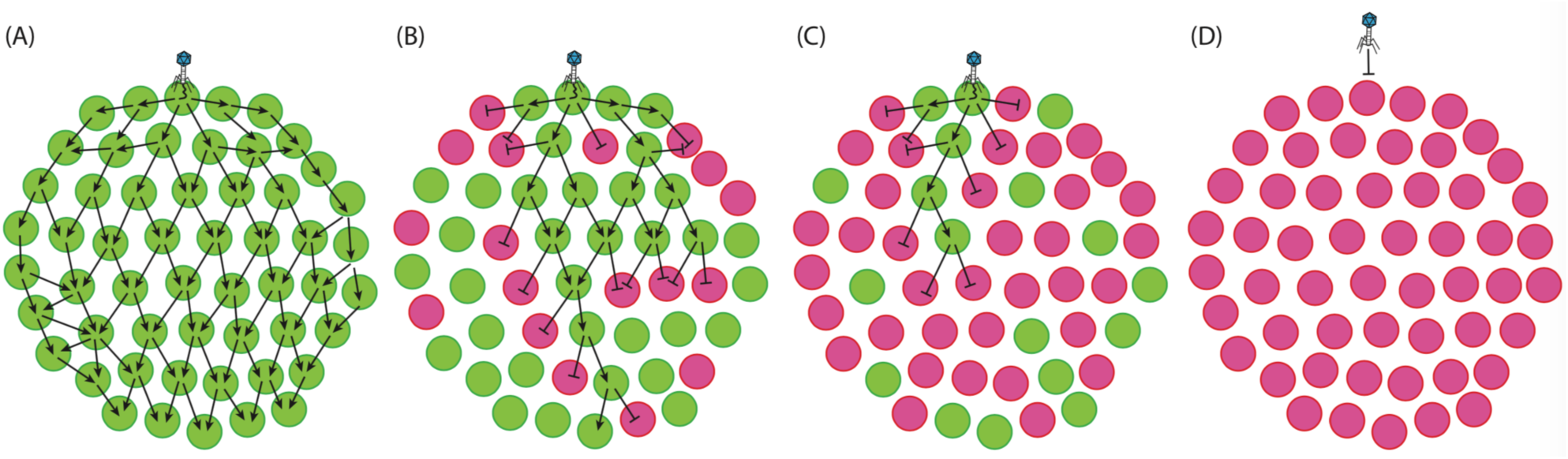
Phase variation of phage resistance genes generates ‘bacterial herd-immunity’ in bacterial meta-populations. Phase variation of a phage-receptor can generate heterogeneous bacterial meta-populations containing a mix of sub-populations that are either susceptible or resistant to phage infection. High proportions of resistant sub-populations can both hinder phage spread and protect susceptible populations from phage attack. In this figure, we show the routes of phage dissemination through four different bacterial meta-populations. Green circles represent phage-sensitive populations while pink circles are phage-resistant populations i.e. *lic2A* ON and OFF populations in our experimental system. Lines with arrowheads show infection events leading to successful phage replication, while lines with barheads represent directions where phage-resistant populations act as a barricade blocking phage spread. Regions free of arrows are those free of phage. Phage spread is shown for: (A) a 100% phage-sensitive (*lic2A* ON) sub-population; (B) 66% phage-sensitive (*lic2A* ON) sub-populations; (C) 66% phage-resistant (*lic2A* OFF) sub-populations; and (D) 100% phage-resistant (*lic2A* OFF) sub-populations.

PV of phage receptors or RM systems is an established phenomenon observed across numerous species and occurring by multiple mechanism. Repeat-mediated switching due to hypermutation of polyG tracts in phage receptor genes of *C. jejuni* strain NCTC11168 abrogates infection by phage F336 (19). Similar polyG hypermutation controls the phage growth limitation system of *Streptomyces coelicor* A3(2) and confers resistance to infection with phage φC31 (33). Epigenetic PV of Ag43 in *E. coli* or the gtr^P22^ operons that control O-antigen modification in *Salmonella* are known or proposed to modulate phage infection (34, 35). High frequency, but not hypermutable, mutations in short polyG or polyA tracts produce resistance to phages in both *Bordetella pertussis* (36) and *Vibrio cholerae* (17). Although these mechanisms vary in the rates of generation of phage resistant variants and in the strength of phage resistance (e.g. high for receptor PV and low for RM PV), they all have the potential to generate spatially-structured populations and hence herd immunity to phage infection.

Additionally, there is potential for evolution of the herd immunity state. PV rates can evolve through changes in the mutable mechanism. Thus repeat-mediated PV rates increase as a function of tract length. We anticipate that frequent exposure to phage would select for a greater capacity to form heterogeneous meta-populations through secondary selection for an increase in mutability of the phage receptor.

### Evidence for immune-driven selection of lic2A ON expression states

One rationale for the existence of PV-driven herd immunity is that protection of the phage-sensitive state is required because this state is advantageous under certain circumstances. For *lic2A*, the ON state in *H. influenzae* strain Rd is phage-sensitive and this states leads to extension of the LOS side-chain from the third heptose with a single galactose, a digalactose or more complex sugars. In *in vitro* studies, *lic2A*-dependent epitopes aid in survival against human immune responses (16), with the LOS extensions associated with *lic2A* expression encoding epitopes also present on the human P blood group antigens (37). Although human volunteer studies of colonisation with *H. influenzae* have found that expression of *lic2A* was not essential for human nasopharyngeal colonisation (26), the *lic2A* ON state has been associated with disease states, including non-typeable *H. influenzae* pneumonia (24). We performed an analysis of 104 *H. influenzae* genome sequences and found that *lic2A* is in an ON state in ∼63% of all isolates and is predominantly in the ON state for multiple disease conditions, suggesting selection for expression of *lic2A* occurs across a wide range of niches for *H. influenzae* (Fig. S5).

These observations are consistent with a scenario where phage drive evolution of hypermutability rates as the *lic2A* gene oscillates between selection for/against the immune-resistant/phage-sensitive and immune-sensitive/phage-resistant states. A caveat to these conclusions is phage-specificity as phage HP1c1 may be specific to extension of the third heptose of *H. influenzae* LOS whereas Lic2A can, in the appropriate genetic context, generate extensions from any of the three heptoses of the LOS inner core. Further work is required to determine whether HP1c1 is specific for extension from the third heptose and if other phages can target this epitope in *H. influenzae* strains where extension is from the first or second heptose.

### Transmission and the ‘phage loss’ phenomenon

Our mathematical model of herd immunity indicates that phage spread is interdependent on population structure and the rate of phage loss from the environment by dilution. Thus, when dilution rates are high, phage extinction events are frequent despite high levels of sensitivity within the bacterial population. While natural rates of phage loss from respiratory environments is unknown, a number of factors have the capacity to impact phage loss, such as humidity, salinity and immune responses (38–41). For phage infections of human commensals or pathogens, such as *H. influenzae*, more extreme environmental selection pressures will apply as phages are transmitted between carriers of target bacterial species.

There are two potential extreme scenarios where the herd immunity model is applicable and ‘phage loss’ is either low or high. These scenarios are elaborated for *H. influenzae* but are relevant to other bacterial commensals/pathogenic bacteria. Firstly, colonisation of asymptomatic carriers by *H. influenzae* is likely to involve a series of microcolonies, a meta-population, distributed across the nasopharyngeal surface rather than one continuous population. Phage will therefore have to transmit between microcolonies thereby imposing a low potential for phage loss such that only high numbers of phage-resistant populations will provide protection for any phage-sensitive microcolonies. A second scenario is a population of *H. influenzae* carriers, in this case phage transmission between carriers is likely to result in significant phage loss and hence low numbers of carriers colonized by phage-resistant populations could prevent phage spread to all carriers. These scenarios illustrate the central role of transmission in shaping bacterial herd immunity and in the impact of this fitness trait on localised hypermutation of the phage receptor.

### Bacterial herd immunity could impact on phage-driven evolution

We observed that phage densities were highly variable across spatially-structured bacterial populations with some regions exhibiting a complete absence of phage (Fig. 6). This imposition of spatially-discrete levels of phage selection could select for alternative adaptive traits within the bacterial host. In studies of *Caulobacter crescentus*, low phage selection led to isolation of >200 phage resistance mechanisms, while only ∼60 distinct resistance forms were isolated during high phage selection (42). Thus, bacterial herd-immunity may prevent uniformity in phage selection pressures across bacterial meta-populations leading to evolution of distinct phage-resistance mechanisms within a single clonal lineage.

In summary, our demonstration of a hypermutable locus retarding spread of an infectious agent within a prokaryotic meta-population suggests that the herd immunity phenomenon may be applicable to a wide variety of interacting biological organisms and have deep evolutionary roots. Our conceptual framework could be utilised to explore whether somatic hypermutation, a key example of localised hypermutation in eukaryotes, evolved through selection for sub-population heterogeneity linked to pathogen resistance.

## MATERIALS AND METHODS

### Phage and bacterial strains used in this study

Phage HP1c1, and the *lic2A* ON (Rd 30S) and OFF (Rd 30R) phase variants of *H. influenzae* were obtained from A. Piekarowicz (University of Warsaw, Poland). Phage HP1c1 is maintained in the lysogenic state within *H. influenzae* RM118-L. *H. influenzae* strains were cultured overnight at 37°C on 1% BHI agar supplemented with 10% Leventhal’s supplement and 2 µg/mL nicotinamide adenine dinucleotide (NAD) or in 10 mL BHI broth supplemented with 2 µg/mL NAD and 10 µg/mL hemin (sBHI).

### Phage HP1c1 stocks

Mitomycin C was added to a concentration of 300 ng/mL to an OD_600_ 0.1 culture of RM118-L in 10 ml of sBHI. After incubation for 8 hours, the culture was centrifuged (4,946x*g*, 4°C, 10 minutes) and then the supernatant was passed through a 0.22 µm syringe filter to obtain a phage suspension, which was stored at 4°C.

High titer phage stocks were propagated in *H. influenzae* Rd 30S by adding 100 µL of induced phage to mid-log phase cultures diluted to an OD_600_ of 0.01 in 10 ml sBHI and incubating for 8 hours. Cultures were processed as described above.

### Determination of phage titers

Phage titers were determined using the small-drop plating assay (43). Briefly, 150 µl of a mid-log phase Rd 30S culture, OD_600_ 0.1, was added to 3 mL of 0.3 % BHI agar (supplemented with 2 µg/mL NAD and 10 µg/mL hemin), mixed by inversion, and poured onto 1% BHI agar plates supplemented with 10% Leventhal’s media and 2 µg/mL NAD. Ten-fold serial dilutions were spotted in triplicate 10 µl drops onto the soft agar (the minimum detection threshold is 33.3 PFU/mL).

### Oscillating prey assay

Two phase variants, namely, Rd 30S (*lic2A* ON, phage sensitive variant) and Rd 30R (*lic2A* OFF, phage resistant variant) were utilized to generate the following three-repeat host population-cycling patterns: S100, ON-ON-ON; S66, ON-ON-OFF; S50, ON-OFF-ON; R50, OFF-ON-OFF; R66, OFF-OFF-ON; R100, OFF-OFF-OFF. *H. influenzae* Rd 30S and Rd 30R were sub-cultured as described above in 20 mL of sBHI to an OD_600_ of 0.1. A 5 mL aliquot of the relevant strain was inoculated with HP1c1 at MOI ∼0.01. Cultures were adjusted to 6 mL, mixed by inversion and 1 mL was removed for filtration and determination of the T = 0 phage titre. The remaining 5 mL was incubated for 50 minutes (i.e. one viral replication cycle) at 37°C with shaking and then filtered for phage quantification. Subsequent cycles were initiated by transferring 600 µl of filtrate to a fresh 5 mL culture of relevant *lic2A* phase variant. Five phage transfers were conducted per day with phage-containing filtrates being stored overnight at 4°C.

### Fixed-area oscillating prey assay

Six cycling frequencies were tested (Fig. 5). Allocation of each phase variant (i.e. Rd 30S or Rd 30R) to specific wells was determined by numbering wells from 1 to 631 followed by randomization of these numbers into two sets using R (see Fig. S3). One phase variant was added to the first set of numbered wells and the other phase variant to the second set.

Rd 30S and Rd 30R were sub-cultured to OD_600_ 0.1 and then 250 µL of appropriate culture was added to the starter well. Phage HP1c1 was added at a final concentration of ∼1 × 10^6^ PFU/mL to this well and the volume was adjusted to 300 µL with fresh sBHI broth. After mixing by tituration, 100 µL was removed for phage titration. The plate was incubated at 37°C with shaking for 70 minutes (a longer incubation time was required in this miniaturised oscillating prey assay for completion of one phage replication cycle [data not shown]). Following incubation, the plate was centrifuged at 1500 × *g* for 4 minutes to pellet bacterial cells and then 20 µl of supernatant was transferred to surrounding wells (see Fig. S4). Remaining supernatant was harvested for phage titration. Newly-inoculated wells received 167 µl of a fresh OD_600_ 0.1 culture of either Rd 30S or Rd 30R, depending on the cycling pattern, followed by repetition of all previously described steps. Ten cycles were performed in the left, right, and downward directions, and 20 cycles in the upward direction (see Fig. S3 and S4). Five transfers and cycles were conducted each day with the 96-well plates being wrapped in paraffin film and stored at 4°C overnight. This assay was conducted once for each population structure.

### Survey of Lic2A expression state in multiple *H. influenzae* genomes

A set of 126 genomes available in GenBank as of 06-09-2016 were screened with blast2seq for the presence of genes homologous to the *lic2A* gene of *H. influenzae* strain Rd KW20. *Lic2A* homologues were found in 121 of the 126 genomes but only 104 could be analysed due to incomplete sequence coverage in 17 genomes. The ON state was identified by the presence of a full length amino acid product (∼300 amino acids). The 5’CAAT repeat numbers were determined by visual inspection of aligned sequences. All metadata, where available, was collated from the GenBank entry for each genome, or references associated with each strain.

### Mathematical model and simulations

We describe bacteria-phage interaction using the conventional modelling approach (45, 46). Dynamics of bacteria and phage densities in each experimental cycle (transfer) of number *n* within time *T*_*0*_=40min (i.e. before the start of mass replication of phages) is given by

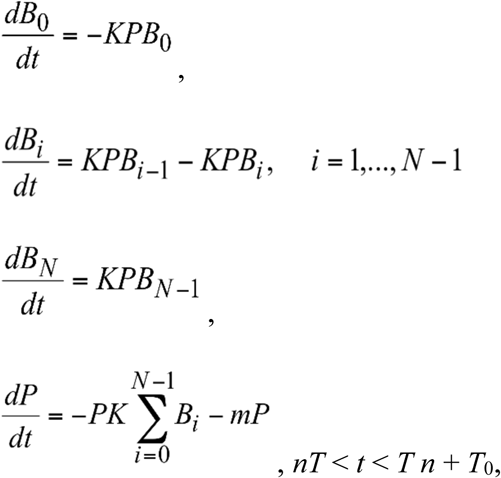

where *B*_*0*_ is the number/density of phage-free bacteria; *B*_*i*_ is the number/density of bacteria with *i* phage attachments; *P* is the number/density of free phages. The maximal number of phage attachments for an individual bacterial cell is given by *N.* In the model, we assume that the injection rate is very fast (i.e. instantaneous), so that attachment of one phage immediately results in an infection.

For simplicity, we assume that there is no bacterial growth. We also assume that all phage attachments occurring within the first 10 minutes lead to replication (each phage produces *b* new phages) whereas later attachments result in phage loss without replication. We neglect binding of newly replicated phages within the last Δ= 10 min of each cycle. The other model parameters are: *K*-, phage adsorption constant (note that this constant is assumed to be independent from the number of bound phages and that *K*=0 for phage-resistant bacteria); *m*, natural mortality of phages; *b*, phage burst size.

At the start of each experimental cycle (i.e. just after dilution) all bacteria are phage-free and their number is always equal to *B*_*S*_, in other words,

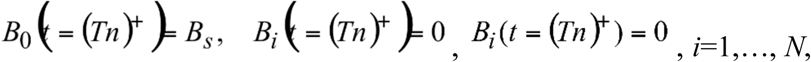

Here the symbol ‘*+*’ denotes the time just after the *n*^th^ dilution, i.e. just prior to the cycle (*n*+1); the symbol ‘−’ denotes the time prior to the *n*^th^ dilution, i.e. at the very end of cycle *n*.

The phage density is obtained from the final density in each cycle multiplied by the dilution coefficient *C*_*n*_ in cycle #*n* (this value can vary between experiments).

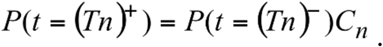

The density of phages just prior to dilution *n* (i.e. at the end of cycle #*n*) is determined by

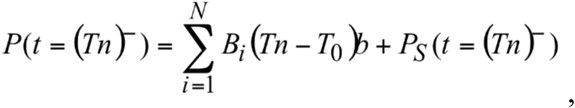

where *P*_*S*_ are non-attached phages that survived to the end of the cycle; i.e. the phage number at the cycle end, prior to dilution, is given by the number of infected bacteria at time *T*−*T*_*0*_ multiplied by the burst size *b* plus the number of surviving phages *P*_*s*_.

Susceptible bacteria are characterized by *K*>0, whereas for resistant bacteria we have *K*=0. The value of *K* is kept constant across each cycle of 50 minutes.

Model parameters and verification were derived from experimental settings or findings. Direct observations indicated that *B*_*S*_=1.75±0.25*10^8^ cell/ml, *T*=50 min, and *T*_*0*_=40 min. The adsorption constant of *K*=7±3*10^−10^ mL/(cell min) was estimated from an adsorption assay (Fig. S1). Other parameters were estimated directly from the oscillating prey assay by model fitting (see Fig. S2): *b*=42±5 (burst size); and *m*=0.006±0.003 1/min. The parameter *N* had a minor effect in our computation and hence we utilised *N* =20 in all subsequent models.

Further simulations (e.g. Fig. 3c) considered the phage-bacterial dynamics across 105 cycles. Variations in the dilution rate *C* were simulated by changing the value of C according to *C*=*C*\*(1+ε), where ε is a normally distributed random variable with a mean of 0. and variance of 0.3^2^. In each cycle, *K* was randomly switched from *K*= 7*10^−10^ (susceptible bacteria) to *K*=0 (resistant bacteria). The frequency of switching was determined by the probability *p*, which gives the probability of encountering susceptible bacteria. Examples simulations, shown in Fig. 3a and 3b, were constructed for *C*=0.1, *p*=0.75 (A) *p*=0.85 (B). Fig. 3c was obtained by repeating simulations across 105 for variable parameters *C* and *p*. Numerical simulation was based on the standard Runge-Kutta integration method of order 4 using MATLAB software. When phage density dropped to or below the low value threshold of *P*_*0*_=100, we considered that this was equivalent to *P*=0 and stopped further simulations. The initial density of phages at time *t*=0 was 4.28*10^7^ PFU/mL.

## Supplemental material

Fig. S1 Parameter setting experiments for mathematical modelling.

Fig. S2 Fit of mathematical model to experimental data.

Fig. S3 Distribution of wells receiving a *lic2A* ON or *lic2A* OFF phase variant of *H. influenzae* strain Rd for the 50 % ON (panel A) and 66 % ON (panel B) population structures.

Fig. S4 Sampling and transfer regimes for testing phage expansion over a fixed area.

Text S1. Modelling the dynamics of PV of the *lic2A* gene of *H. influenzae*

Fig. S5 Putative ON/OFF state of the *lic2A* gene from 104 *H. influenzae* strains.

