## Supplementary Materials for "Oscillating bacterial expression states generate herd immunity to viral infection"

**Fig. S1 Parameter setting experiments for mathematical modelling.**

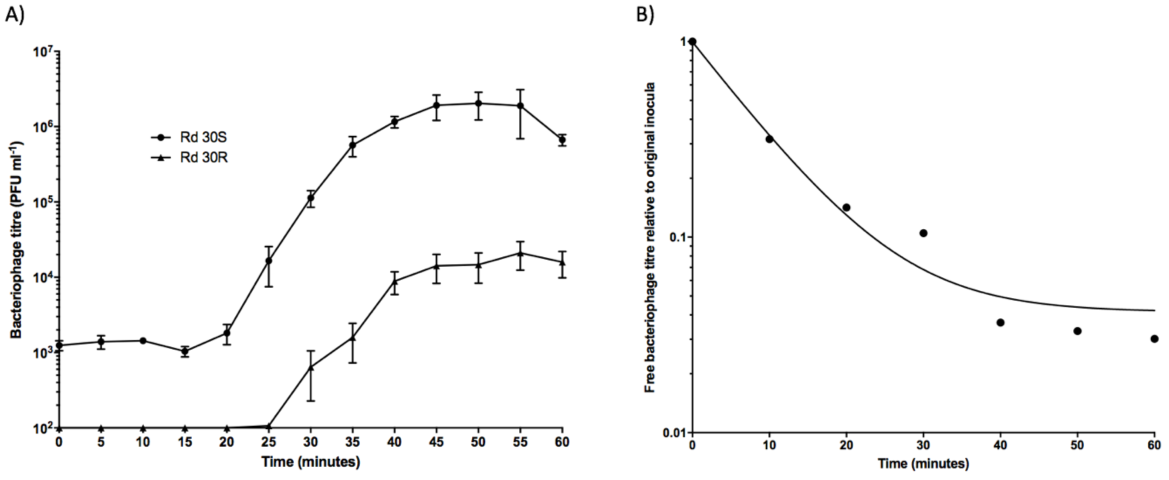


Panel (A) shows a phage one step growth curve used to determine replication time of phage HP1c1 and the burst size of the phage in both phage sensitive and resistant populations. Note that phage replication is still observed in phage resistance populations due to a low level of phage sensitive phase variants where the *lic2A* gene has switched back to the phage sensitive ON state. Figure (B) shows phage adsorption assays for phage HP1c1. The phage was incubated with the phage sensitive *H. influenzae* strain (Rd 30S) and the phage remaining in the media was determined at defined intervals, resulting in a measure of the rate of phage adsorption to host cells. The estimated value of the adsorption constant of phages is $K$=7±3x$10^{-10}$ ml/(cell min) and was estimated from the linear portion of the curve.

Methods. **One-step growth curve of phage HP1c1**. Overnight culture of either Rd 30S or Rd 30R were sub-cultured to $OD_{600}$ 0.1 (as above) in 40 ml sBHI with phage HP1c1 then added to a final concentration of 1x$10^5$ PFU mL$^{-1}$ (MOI 0.001), mixed, with 1 ml then removed and passed through a 0.22 µm filter to determine the initial phage concentration. The remaining suspension was then incubated at 37°C for 10 minutes to allow phage to bind cells, after which the culture was centrifuged at 4,696 x *g* for 5 minutes at 4°C. The supernatant was discarded and the bacteria/bound phage pellet was resuspended in 40 ml fresh sBHI pre-chilled to 4°C. Centrifugation and washing in chilled broth was repeated again. After this, the bacteria/phage pellet was resuspended in 40 ml room temperature sBHI and aliquoted into 13x 2 ml aliquots, with these then incubated at 37°C, shaken at 100 rpm . Aliquots were removed for phage titration at 5 min intervals over 60 minutes.

**Phage HP1c1 adsorption assay**. HP1c1 adsorption rate was measured as previously described (44). Rd 30S was sub-cultured in 10 ml sBHI to $OD_{600}$ 0.1 as described above, with 1 ml then removed and used to determine bacterial density by spreading 100 µl of ten-fold serial dilutions onto BHI 1% agar supplemented with 10% Leventhal's supplement and 2 µg mL$^{-1}$ NAD and incubated overnight at 37°C. To the remaining 9 ml culture, 2 µl of 30 mg mL$^{-1}$ chloramphenicol was added preventing phage replication during the assay. The culture was then incubated at 37°C, agitated at 100 rpm, for 5 minutes to allow the suspension to reach 37°C before phage addition. After which 1 ml of prewarmed HP1c1 was added to a final concentration of ~1x$10^7$ PFU mL$^{-1}$. At 10 minute intervals 50 µl of the suspension was removed and added to ice-chilled tubes containing 950 µl sBHI broth plus three drops of chloroform with these tubes then briefly vortexed, and placed back on ice until completion of the experiment. Phage titre was determined for each tube by adding 100 µl of the phage suspension to 150 µl of $OD_{600}$ 0.1 Rd 30S, followed by addition of 3 ml BHI 0.3% agar supplemented with hemin and NAD. This was then mixed by inversion and spread onto BHI 1% agar plate supplemented as above, allowed to set and then incubated overnight at 37°C.

**Fig. S2 Fit of mathematical model to experimental data.**

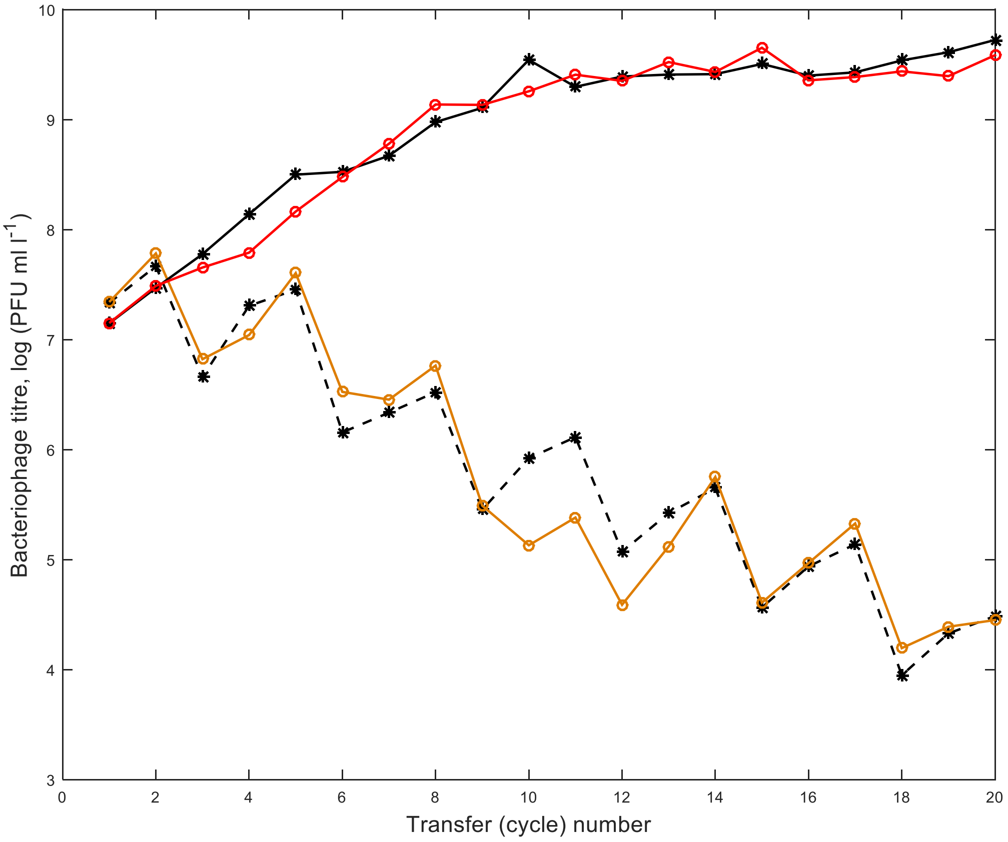


The figure shows the fit of our mathematical model to the oscillating assay data (we plot the phage density at the end of each cycle) for the cases where all bacteria are susceptible (S100%) and where 66% of bacteria are susceptible (S66%). Coloured lines correspond to the experimental data; black lines show our model fit. The estimate of the model parameters are $b$=42±5 (burst size) and $m$=0.006±0.0031/min (natural mortality). The other model parameters are given by $K=6\text{x}10^{-10}$ ml/(cell min); $T$=50min; $T_0$=40 min; $B_S=1.75\text{x}10^8$ cell/ml; $N$=20. The values of dilution coefficients $C_n$ were calculated directly from experimental data by taking the ratio of the phage densities before and after dilution.

**Fig. S3 Distribution of wells receiving a *lic2A* ON or *lic2A* OFF phase variant of *H. influenzae* strain Rd for the 50 % ON (A) and 66 % ON (B) population structures.**

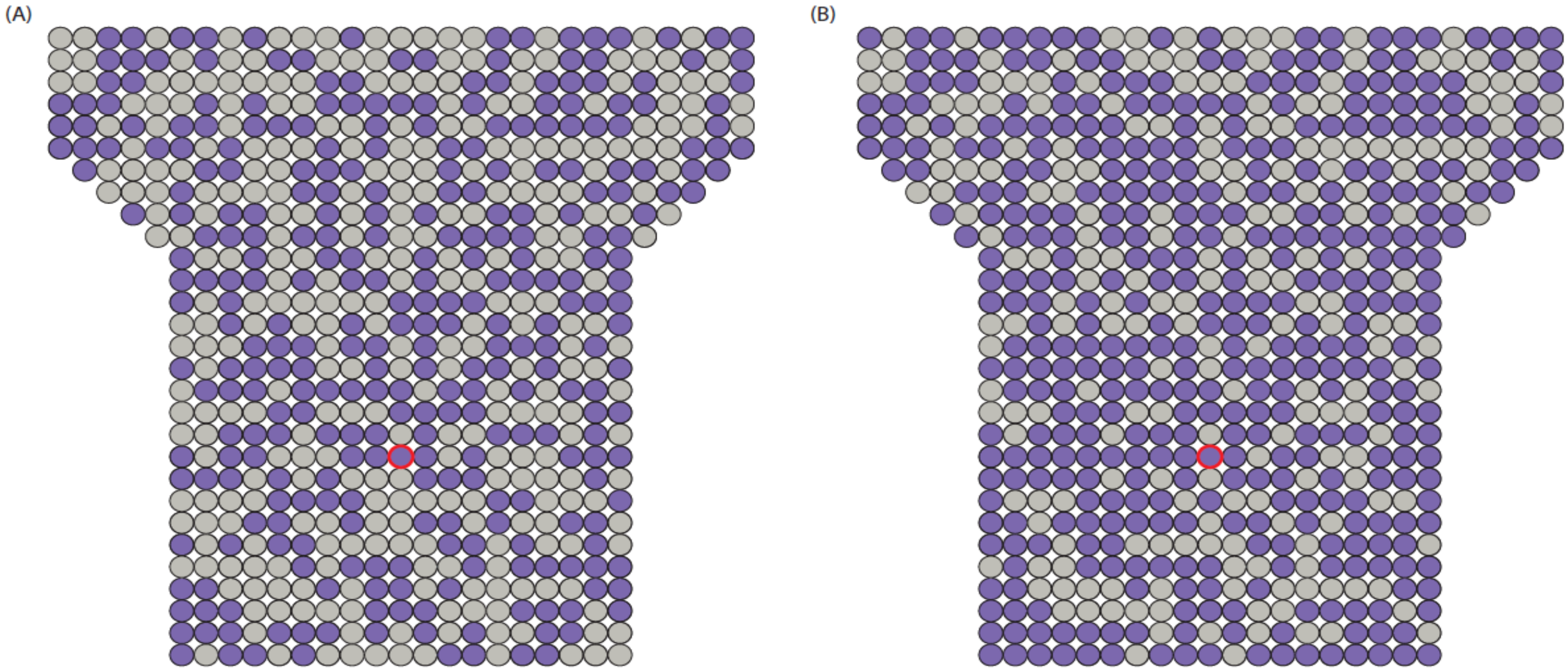


The phase variants of *H. influenzae* strain Rd were grown to mid log phase in BHI broth and this culture was utilised as the inoculum for each well of the spatial assay. Note that these inocula will contain a population in which ~99.9% of the cells are in the expected expression state and ~0.1% in the alternate state due to PV during growth of the culture. In both examples, the filled purple circles represent wells inoculated with a *lic2A* ON population, while grey filled circles represent the *lic2A* OFF populations. For the 50 % OFF and 66 % OFF populations, the fill patterns for the wells were reversed**.**

**Fig. S4 Sampling and transfer regimes for testing phage expansion over a fixed area.**

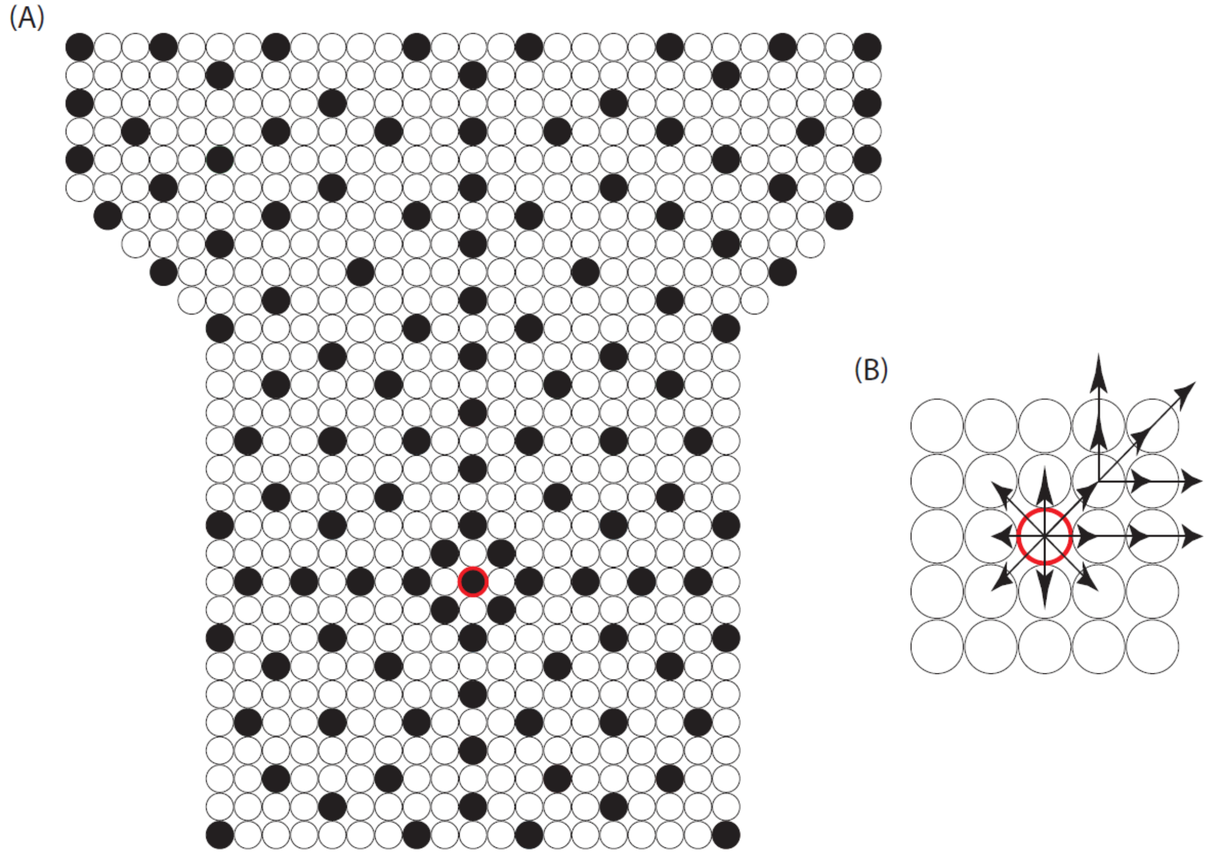


Panel (A) shows the structure of the fixed area over which phage expansion was measured. Each circle represents one well in a 96-well microtitre plate. Filled black circles indicate the wells from which samples were collected from measurement of phage titre. Red circle perimeter shows the well that received the inoculum. Panel (B) shows the pattern of transfer, emanating from the initial well that was inoculated with the HP1c phage, for each sequential cycle of phage infection.

**Fig. S5 Putative ON/OFF state of the *lic2A* gene from 104 *H. influenzae* strains.**

**(A)**

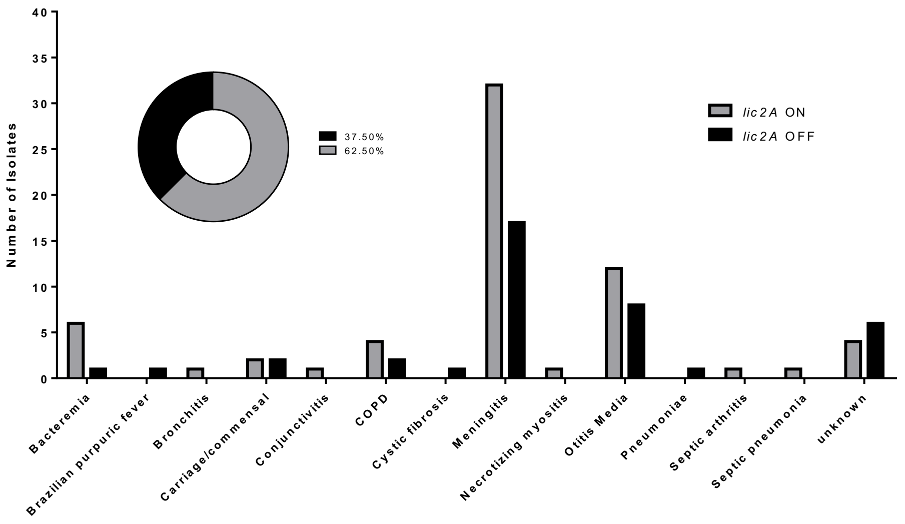


**(B)**

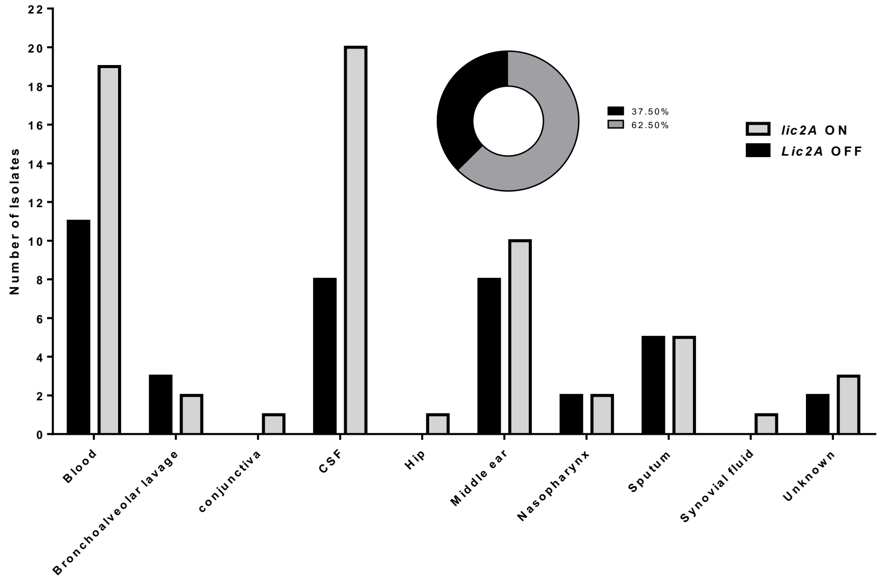


Genome sequences were extracted from the genbank database and searched for the presence of an intact *lic2A* gene. The number of repeats in each *lic2A* were determined and utilised for derivation of the expression state. Panel A is a bar graph of the numbers of isolates with ON (black bars) or OFF (grey bars) *lic2A* expression states for the different clinical states associated with each isolate. Panel B depicts the same data but utilising the site of isolation for each isolate.

**Text S1. Modelling the Dynamics of Phase Variation of the *lic2A* Gene of *H. influenzae***

The *lic2A* gene of *H. influenzae* is subject to phase variation (on/off switching) due to mutations in a 5'CAAT repeat tract present within the reading frame. In *H. influenzae* strain Rd, the ON state of the gene correlates with sensitivity (S) to infection with bacteriophage HP1c1 while the OFF state is associated with resistance (R) to infection. In the absence of phage, the relative proportions of the R and S states will reach a steady state determined by the rates of switching i.e. ON-to-OFF and OFF-to-ON. However, the ON state of the gene is associated with resistance to serum mediated-killing of *H. influenzae*, so that in the absence of phage and the presence of serum there will be selection for the ON state of the gene. The degree of selection by serum components acting on *H. influenzae* occurring within upper respiratory tract during normal host colonisation is unknown but is assumed to be weak. The relative proportions of the ON and OFF states of the *lic2A* gene will therefore be determined by the combination of the switching rates and the level of selection. The aim of this model was therefore to determine the relative proportions of the S and R states for phage infection for a range of different PV rates and selection pressures for S, the phage-sensitive, serum-resistance state, ON state of the *lic2A* gene.

Switching rates for the *lic2A* gene are taken from previously established switching rates (1) where it was estimated that ON-to-OFF switching of this gene occurred at a rate of $1.89*10^{-4}$ and OFF-to-ON switching at $1.13*10^{-4}$. The carrying capacity for the upper respiratory tract is estimated at $10^9$ cfu. The bacterial population is assumed to turnover every hour with a generation time of one hour per division.

In the model, *S* and *R* are the densities of non-resistant (susceptible) and resistant bacteria (cell/l), respectively. We assume for simplicity that the length of the cell life cycle is constant. The mortality rate of bacteria is suggested to be density dependent. The densities of *S* and *R* at time t (measured in hours) will be given by

$$S(t) = b[S(t-1)(1-\alpha) + R(t-1)\beta]\exp(-b(S(t-1) + R(t-1))/K),$$

$$R(t) = b[S(t-1)\alpha + R(t-1)(1-\beta)]\exp(-b(S(t-1) + R(t-1))/K)m,$$

where $b$=2 is the number of cells obtained after each division; $K$=$10^9$ is the carrying capacity (the total maximal number of bacteria that the environment can carry); $\alpha$ is the probability of transition $S \rightarrow R$ per cell division, $\beta$ is the probability of transition $R \rightarrow S$ per cell division. The parameter m shows a drop in the fitness of *R* with respect to that of *S* due to extra mortality of *R* (we call it selection). We consider that $0< 1- m <<1$, i.e. m is smaller but rather close to unity. In other words, the fitness coefficients of resistant and non-resistant bacteria are close to each other.

We start computation from the condition that *S*(0)=10 cell/l; *R*(0)=0.

One can see from Fig. 5 that for the same ratio between the probabilities of switching the final proportion of R $\{R/(R+S)\}$ in a population would strongly depend on fitness. The actual predicted switching rates would result in 1% and 30% of the cells being in the phage resistant state for weak (m=0.99) versus no selection respectively. For fast mutation rates (higher rates of $\alpha$, $\beta$), low level selection (m=0.999) would result in 9% phage resistant variants, however low switching rates combined with low levels of selection would drive the levels of this phage-resistant state to ~0.1% (m=0.999).

We can also calculate the equilibrium ratio of resistant bacteria in the system $R/(R+S)$ analytically using the condition that $S(t+1)=S(t)=S$; $R(t+1)=R(t)=R$. We obtain

$$S/R=\frac{S(1-\alpha)+R\beta}{S\alpha+R(1-\beta)}\frac{1}{m}=\frac{S(1-\alpha)/R+\beta}{S\alpha/R+(1-\beta)}\frac{1}{m},$$

We denote $\varepsilon=S/R$ and obtain.

$$\varepsilon=\frac{\varepsilon(1-\alpha)+\beta}{\varepsilon\alpha+(1-\beta)}\frac{1}{m}$$

For the value of $\varepsilon$ (epsilon), we arrive to the following quadratic equation

$$\varepsilon^2\alpha m+\varepsilon(m(1-\beta)-1+\alpha)-\beta=0.$$

One can see that the above equation always has a unique positive solution.
The value of $\varepsilon$ and $\eta=R/(R+S)$ are related via $\eta=1/(1+\varepsilon)$.

**Fig. S5 Putative ON/OFF state of the *lic2A* gene from 104 *H. influenzae* strains.**

**(A)**

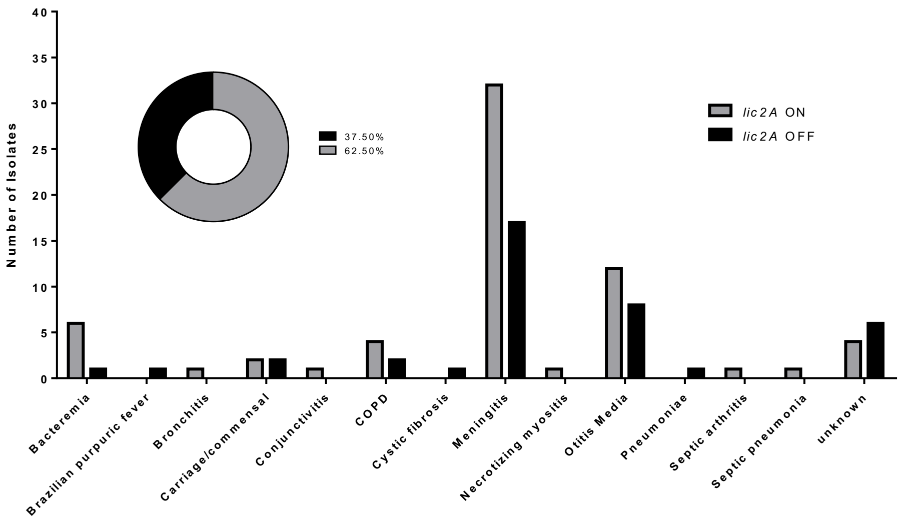


**(B)**

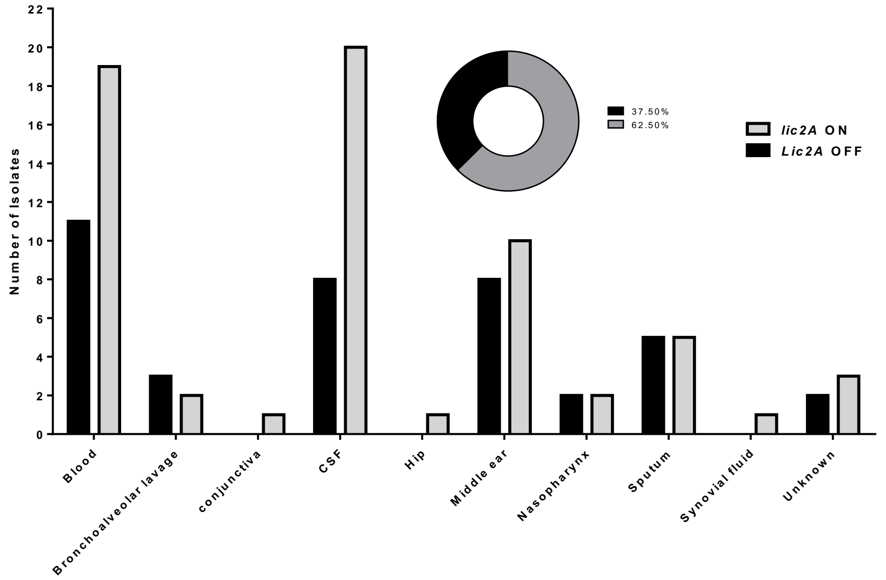


Genome sequences were extracted from the genbank database and searched for the presence of an intact *lic2A* gene. The number of repeats in each *lic2A* were determined and utilised for derivation of the expression state. Panel A is a bar graph of the numbers of isolates with ON (black bars) or OFF (grey bars) *lic2A* expression states for the different clinical states associated with each isolate. Panel B depicts the same data but utilising the site of isolation for each isolate.
